# Single cell RNA sequencing reveals cellular diversity of trisomy 21 retina

**DOI:** 10.1101/614149

**Authors:** Jihong Wu, Xiangning Ding, Fangyuan Hu, Zaoxu Xu, Shenghai Zhang, Langchao Liang, Chaochao Chai, Jixing Zhong, Shiyou Wang, Xiumei Lin, Yin Chen, Qikai Feng, Jiacheng Zhu, Sanjie Jiang, Jun Xia, Wei Li, Ya Gao, Jiankang Li, Jingxuan Zhang, Zhikai He, Shen Xue, Guanglin Guo, Ping Xu, Fengjuan Gao, Dandan Wang, Daowei Zhang, Dongsheng Chen, Xinghuai Sun, Fang Chen

## Abstract

Retina is a crucial tissue for the capturing and processing of light stimulus. Characterization of the retina at single cell level is essential for the understanding of its biological functions. A variety of abnormalities in terms of morphology and function were reported in T21 retina. To evaluate the effects of chromosome aneuploidy on retina development, we characterized single cell transcriptional profiles of a T21 fetus and performed comprehensive bioinformatic analyses. Our data revealed the diversity and heterogeneity of cellular compositions in T21 retina. In total, we identified seven major cell types, and detected several subtypes within each cell type, followed by the detection of corresponding molecular markers including previously reported ones and a series of novel markers. Our analyses identified extensive communication networks between distinct cellular types, among which a few ligand-receptor interactions were associated with the development of retina and immunoregulatory interactions. Taken together, our data provided the first single cell transcriptome profile for human T21 retina which facilitates our understanding on the dosage effects of chromosome 21 on the development of retina.

## Introduction

Retina is a highly specialized neural tissue that contains a variety of neurocyte types that sense light and initiate image processing. Retinal development begins with the establishment of SHH and SIX3 protein-mediated eyeballs, followed by PAX6 and LHX2 proteins to regulate the development of optic nerve vesicles^1^. The vesicles produce three structures: the neural retina, the retinal pigment epithelium, and the rod. The neural retina contains retinal progenitor cells (RPC), which produce seven cell types of the retina^2^. Akina Hoshino, et al. divided the development of the retina into three periods according to RNA-Sequencing. Progenitor cell proliferation and ganglion cell production predominate in the early retina ^3^. The second period is characterized by the appearance of horizontal cells and amacrine cells. At this time, synapse-related genes also showed significant up-regulation. The third period showed the production and differentiation of photoreceptors, bipolar cells and Müller glial cells^4,5,6^.

Trisomy 21, also known as Down syndrome, is the most common birth defect caused by abnormal chromosome dosage with a worldwide incidence reaching 1 in 700^7^. Patients with Down’s syndrome may have clinical manifestations such as congenital heart defects, premature aging, early onset of Alzheimer’s disease and leukemia, as well as retinal developmental abnormality^8^. One of the clinical complications in children with Down syndrome is ophthalmic disease. T21 patients were reported to have abnormalities in neurological pathway^9^. Sharon J. Krinsky-McHale, et al. reported that adults with Down syndrome have significant visual impairment relative to the normals^10^.

Although it has been reported T21 individuals suffered from retina diseases, but the cellular compositions and related molecular regulatory mechanism in T21 retina remains largely unknown. The single cell RNA-sequencing (scRNA-seq) technique has been widely employed to profile the transcriptome of retina in human, mouse, monkey and chicken^11,12,13,14^, which comprehensively characterized transcription profiles of retina cells and attempted to identify the gene regulatory networks of neurogenesis and cell fate specification^15^. To explore the heterogeneity of T21 retina, we performed scRNA-seq analysis on the retina from a fetus at the gestational age of 22 weeks. According to the study in time series in single cell resolution of human retinal development, in the mid-gestational stages (GW14 to GW27), the cell types of retina were specified by retina progenitor cells sequentially. Especially the GW22 was the start point of emergence of rod photoreceptors.^11^ In this study, we examined the gene expression profiles of T21 retinal cells, identified major retina cell types and explored the significant influence of redundancy chromosome 21 in certain cell types. Finally, we trained a machine learning model using random decision forests to make classification of the 21 cells from normal cells.

## Result

### Collection of the 21-trisomy retina tissue and single cell RNA-seq

We collected retinal biopsies from a trisomy 21 donor and dissociated the sample into single-cell suspension without surface marker pre-selection, followed by scRNA-seq (Methods). After sequencing, we obtained the single-cell transcriptome of 3136 cells, with a mean coverage of 115,969 reads per cell (Figure S1b). After filtering, a total of 2866 cells were retained for subsequent analysis (Figure 1).

**Figure 1.**
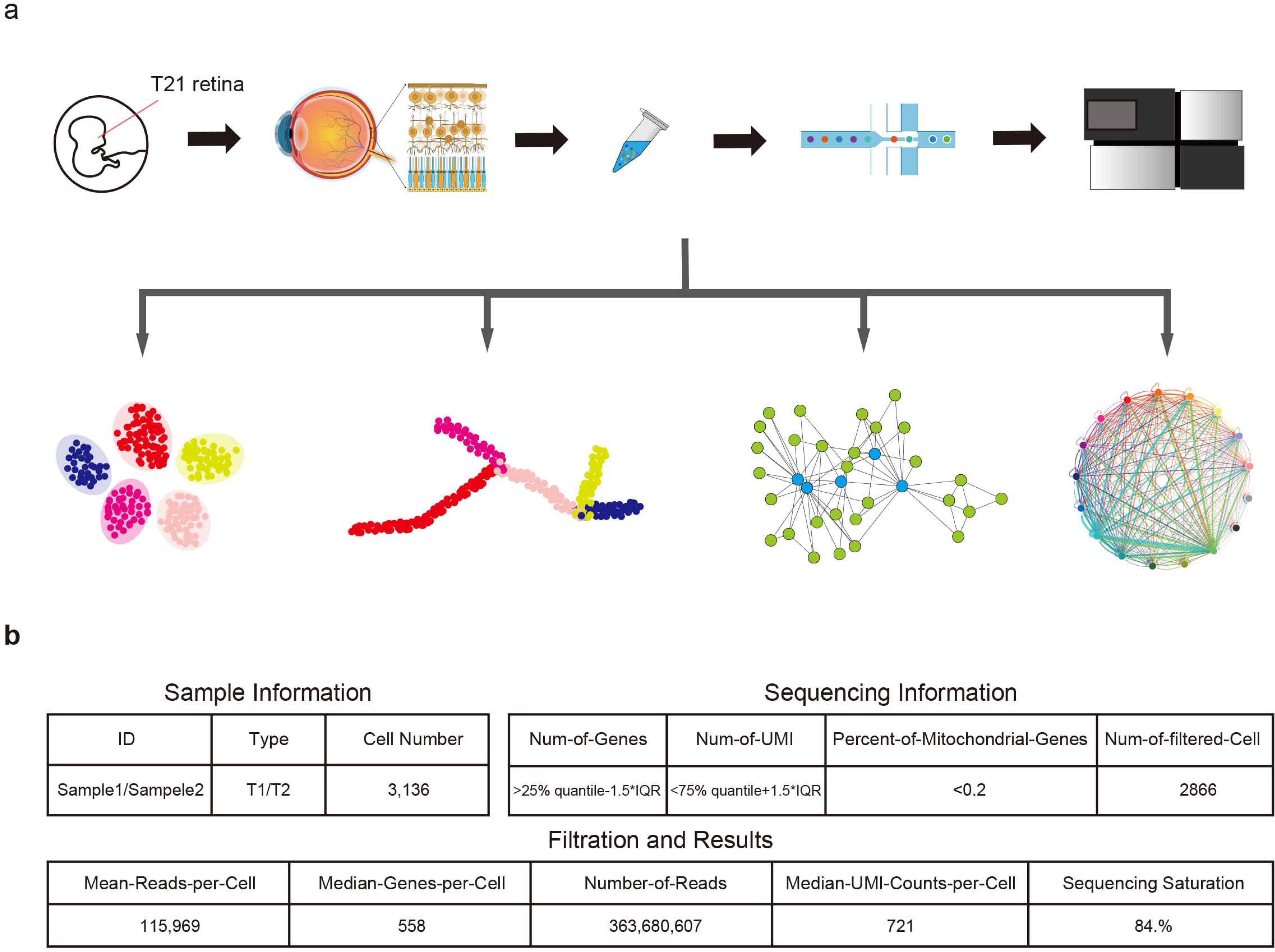
Generation of T21 human fetal retina scRNA-seq data set. a. Schematic representation of the experimental workflow. b. Summary statistics for sequencing information and filtration results of the sample.

### Cellular heterogeneity in retina tissues

Using unsupervised clustering method, 2866 cells were classified into 10 major clusters, corresponding to seven cell types (retinal progenitor cells, bipolar cells, rod photoreceptor, cone photoreceptor cells, retina ganglion cells, Müller glia and astrocytes) (Figure 2a). Briefly, C0, C1, C4, C7&C9, C3&C5&C6 were annotated as retinal cells based on specific expression of previously reported cell types markers and GO term enrichment analysis of cluster specific expressing genes. C0, expressing retina progenitor markers *OTX2*, was defined as retina progenitor cells (RPC). C1 was composed of bipolar cells (BC), according to the expression of *TRPM1*. C4 was considered to be retina ganglia cells due to the enrichment of *RBPMS*. C7 and C9 were identified as cone photoreceptor, because of the specific expression of *LMOD1* ^16^(Figure 2c, d, e, Figure S2).

**Figure 2.**
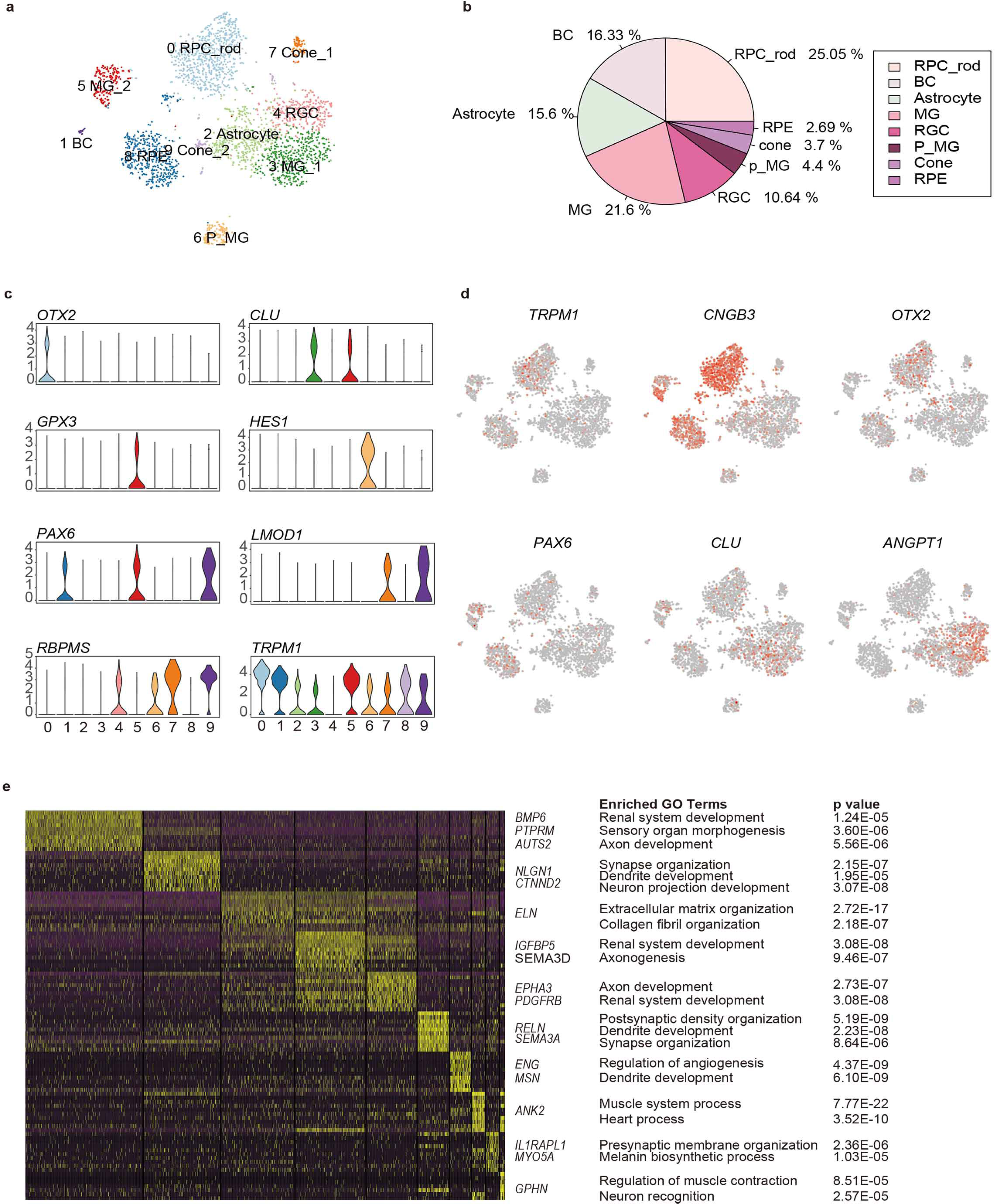
Cellular diversity in retina tissue. a. Clustering of 2866 single-cell expression profiles into 10 retinal cell populations and 2D visualization of single-cell clusters using t-SNE. Each dot represents a single cell, and clusters are color-coded. b. Pie plot of cell types proportion in the trisomy 21 sample. c. Violin plots demonstrating expression levels of marker genes for cell cluster. d. T-SNE maps of retina single-cell data with cell colored based on the expression of marker genes for particular cell types. Gene expression levels are indicated by shades of red. e. A gene expression heatmap showing top differentially expressed gene for each cluster in 10 major clusters. Yellow corresponds to high expression level; purple and black correspond to low expression level. Significant marker genes were listed on the right as well as enriched GO terms and corresponding P values.

C3, C5 and C6 expressed three different Müller glia cell markers respectively, *CLU, GPX3* and *HES1*^17^. C6, displayed characteristics of Müller glia cell with differentiation potential, through the high expression of *HES1* (marker for retina progenitor cells) and the enrichment of development Gene Ontology such as dendrite development. The distinction between C3 and C5 revealed two functional subtypes of Müller glia cells, with C3 being related to renal system development and axonogenesis while C5 specifically expressing *GPX3* and enriching for GO terms associated with synapse organization. Trajectory analysis further confirmed that these three clusters had differential relationship and seemed to represent three differentiation stages of Müller glia cell^11^.

C2 and C8 were annotated as astrocytes and retina pigment cells respectively. C2 was characterized by the high expression of *ANGPT1*. C8, expressing markers of retina pigment cells, such as *TYRP1* and *RMEL*^16^, was defined as retina pigment cells.

To explore the cellular compositions of T21 retina, we calculated the proportion of each retinal cell type. It was found that the most abundant cell type was rod photoreceptors, followed by Müller glia and bipolar cells. Cone photoreceptors and RPE accounted for only 3.7% and 2.69% respectively. We noticed that two major retinal cell types (horizontal cells and amacrine cells) were undetected in our data, probably because of their relatively low proportion. The cell type proportion of the sample (Fig 2b) demonstrated the abnormal constitution of retina neuron compared with normal fetal at the same gestational weeks according to recent researches^11,17^.

### Dissection of retina neuron states transition through developmental trajectory analysis

To dissect the differentiation process among human retina, we first removed RPE and glia cells due to the irrelevance in retinal developmental relationship ^18^ and then merged remaining retina neural cell clusters to create a systematic landscape for the whole cell lineage using Monocle ^19,20,21^. Each cluster was ordered along with the pseudo-time (Figure 3a). To further investigate the mechanism of cell types determination, we interrogated 113 transcription factors which expressed variably along the constructed developmental path. SOX5^22^ specifically expressed at progenitor cells and NFATC4 specifically expressed at cone photoreceptor cells had been shown to regulate the neuron morphogenesis which played positive and negative role respectively (Figure 3c). Previous study reported that POU6F2^23^ could promote stem cells commitment to ganglion cell fate while RORB^24^ showed the capacity to facilitate cone cell development (Figure 3c). The transcription factors (TFs) ontology was enriched to cast the tree of term according to the similarities among their gene memberships (Figure 3c, d). MCODE algorithm was then applied on the network to identify neighborhoods where proteins were densely connected (Figure 3e). We then calculated differentially expressed genes of each branch of development trajectory and clustered the genes according to the expression pattern between branches (Figure 3f). There were two main specific patterns of gene expression. Gene cluster 2 was specifically expressed in RPCs, enriching in neural system development, while gene clusters 1 was specifically expressed in BCs, enriching in inorganic cation transport and synapse localization (Figure 3g, h). To further illuminate the key regulatory elements during retina neural development, we then constructed the regulatory network using nine TFs that not only differentially expressed between branches but showed neurodevelopment related functions. By calculating the co-expression pattern of genes with the selected TFs (Figure S3a) and then extracting the top1000 links showing positive correlation with each TF, we revealed a densely connected network for retinal cell type specification (Figure S3c). Next, we inferred a regulatory network among all differentially expressed TFs based on known interactions collected in the STRING database^25^.

**Figure 3.**
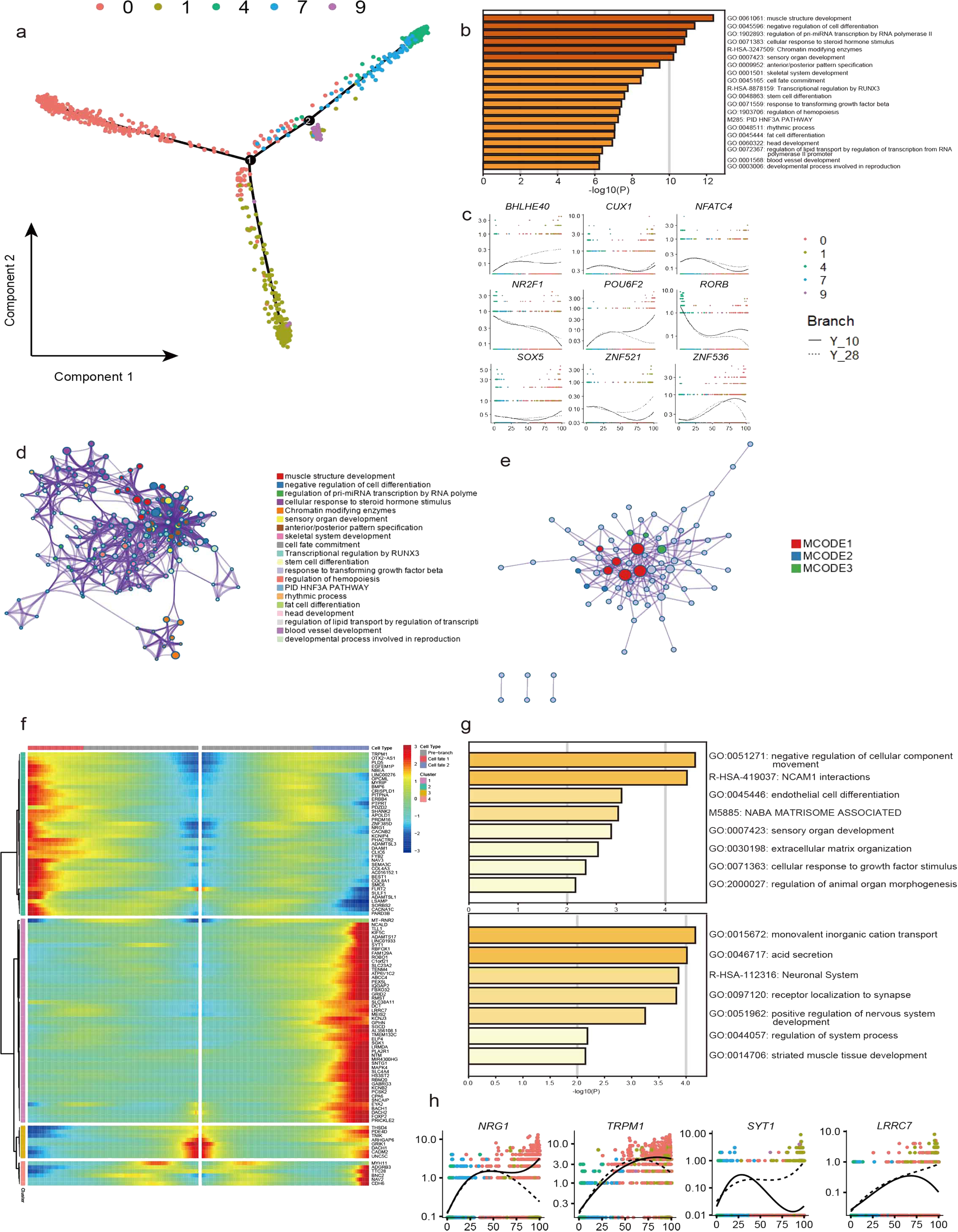
Developmental trajectory analysis of the retina neural cells state transition. a. Single cell trajectories by Monocle analysis showing development of the T21 human fetal retinal neural cells. b. g. Enriched GO terms of variably expressed TFs along the trajectory and variably expressed genes along the branches respectively.. c. h. Expression profiles of selected cell lineage master regulators on developmental trajectory and key genes on branches respectively. d. Accumulated hypergeometric p-values of identified statistically enriched terms were hierarchically clustered into a tree based on Kappa-statistical similarities among their gene memberships. Selected subsets of representative terms from these clusters and converted them into a network layout. e. MCODE algorithm (supplementary note) was then applied to this network to identify neighborhoods where proteins are densely connected. f. Changes for the genes that are significantly branch dependent.

### Identification of master regulators specifying photoreceptor cells development

Next, we reconstructed the developmental trajectories of photoreceptor cells. To infer TFs contributing to cell status transition, we constructed the regulatory network using differentially expressed TFs for retina progenitor cell and photoreceptor cells. We constructed the regulation network of these 13 TFs by analyzing the genes co-expressed with TFs (Figure S4b) and extracted the top1000 links showing positive correlation with each TF. The result suggested that all these 13 TFs demonstrated strong regulatory function and *MYRF, PRDM16, BNC2, HMGA2* and *PBX1* were key regulators of the network since extensive co-regulation with potential target genes was showed. Consistent with our functional annotation result, these key regulators played important roles in the process of retina formation and the light signal transition of photoreceptor (Figure S4c).

### Construction of Müller glia cell differentiation trajectory

To characterize the Müller glia with differentiation potential, we extracted the Müller clusters and constructed the trajectory with unsupervised clustering gene. C6 lay at the root state, consisting of the specifically expressing of developmental markers, while C3 and C5 respectively lay at the two ends of development trajectory. We then compared the TF expression at three states, and found that *MEIS1* was expressed specifically in C5, of which top 1000 target genes enriching Gene Otology of positive regulation of neurological system process and JUND highly expressed in C3 and C5, correlating with neurotransmitter secretion and transport. Interestingly, *CUX1* and *FOXP2* expressed in C3 and C5, whose top 1000 target genes enriching immune regulatory function terms, suggesting the multiple supportive and regulatory function of Müller glia during human retina development (Figure S4 f, g, h, i).

### Widespread cross-cell type and intercellular communication network

Based on the identification of several different Müller cell types and astrocytes, whose intense communication with other retinal cell types has been found in previous study^26^, we further moved on to explore the cross-cluster and intercellular communication network within retina. Müller glia fulfil many crucial roles, supporting neuronal development, survival and information processing^27^. The cellular interactions were inferred using public ligand-receptor database (Methods). The expression patterns of ligand-receptor pairs in the networks revealed dense cross-cluster and intercellular communication networks^28^ (Figure 4a, stable 1). Briefly, in cross-cluster network, 56 ligands and 36 receptors were expressed within C3, and the most frequently interactions were observed in the subtype of Müller glia cell while C6 and C7 were the clusters that received the most interactions. To further explore the roles of Müller glia cells in supporting neural cells, we extracted the communication pairs between Müller glia cells and other retinal neural cells to compare both the shared and cell-type specific interactions (Figure 4c,4e). We found *ITGAV* receptor (expressed in all retinal neural cell types) (Figure 4d,4f), together with ligand *FBN1* to be the most frequent interaction pair in Müller glia signaling network. The *FBN1-ITGAV* pair was shown to be involved in mediating R-G-D-dependent cell adhesion^29^. Likewise, the widely expressed receptor *ROBO1*, presented simultaneously with its cognate counterpart *SLIT2*, holds a crucial role in the regulation of commissural axon pathfinding^30^. Besides, *LRP1* expressed in C4 and *CD74* expressed in C6, might both bind to the APP protein expressed by C3, which are associated with different signaling circuits regarding the formation and function of retina^31^ (supplementary table2). Another observation is that the Müller glia-expressing ligand CTGF, by signaling through *ERBB4* (specifically expressed in C0), could regulate nervous system development according to a previous study^32^. As for the communications between Müller glia cell with all other glia cell types, *DDR2* was found to be specifically expressed in astrocytes and together with *COL3A1* and *COL1A1*, involved in the maintenance of immune homeostasis in retina^33^. The densely connected communication network revealed a global signaling interactions within retinal cells, plenty of which still deserve thorough studies in the future.

**Figure 4.**
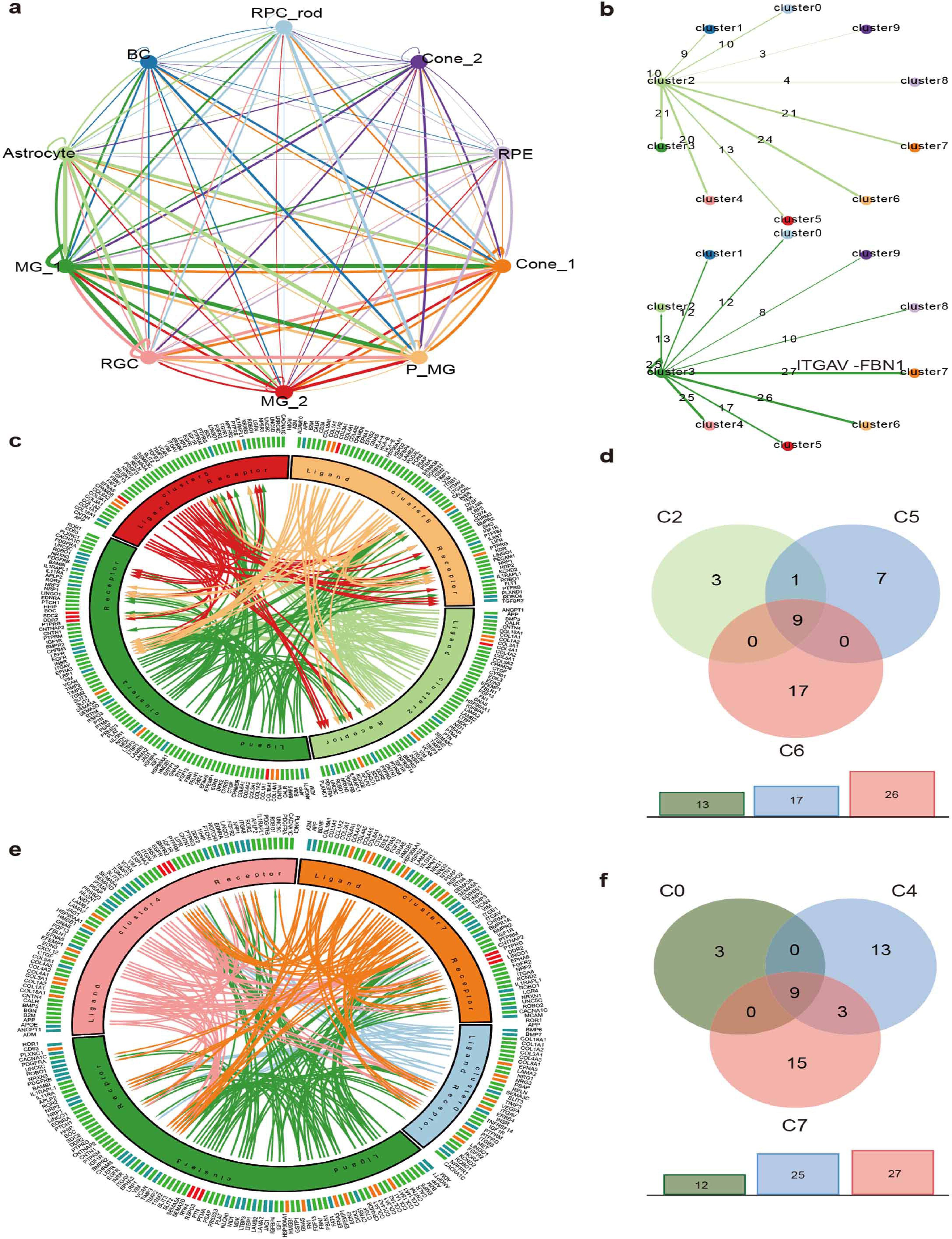
a. Summary of the communication network broadcast by major 10 clusters and those populations expressing cognate receptors primed to receive a signal. b. Detailed view of Müller glial subtype and astrocytes ligands and the paired expressed receptors numbersa. c. e. Visualization of the interactions between Müller glial cells subtype and other glia cells and other retina neural cells respectively. d. f. Comparison with communication pairs of each individual cell type regulated by Müller glial cells subtypes.

### Investigation of the expression profiles of chromosome 21 encoding genes

To study the functional implications of trisomy 21 on retina, we analyzed the expression profiles of genes located on chromosome 21(Figure S6). Next, we performed hypergeometric enrichment test (Methods) and found RPCs(C0) and one subtype of Müller glia cell(C6) were significantly enriched for chromosome 21 encoding genes, indicating the association between the influence of redundant chromosome 21 and the specific cell types (Figure 5a). Also, we found RPCs specifically expressed *CLIC6* and the Müller glia subtypes which has developmental potential specifically expressed *ERG* and *ETS2*, all of which are encoded by chromosome 21.

**Figure 5.**
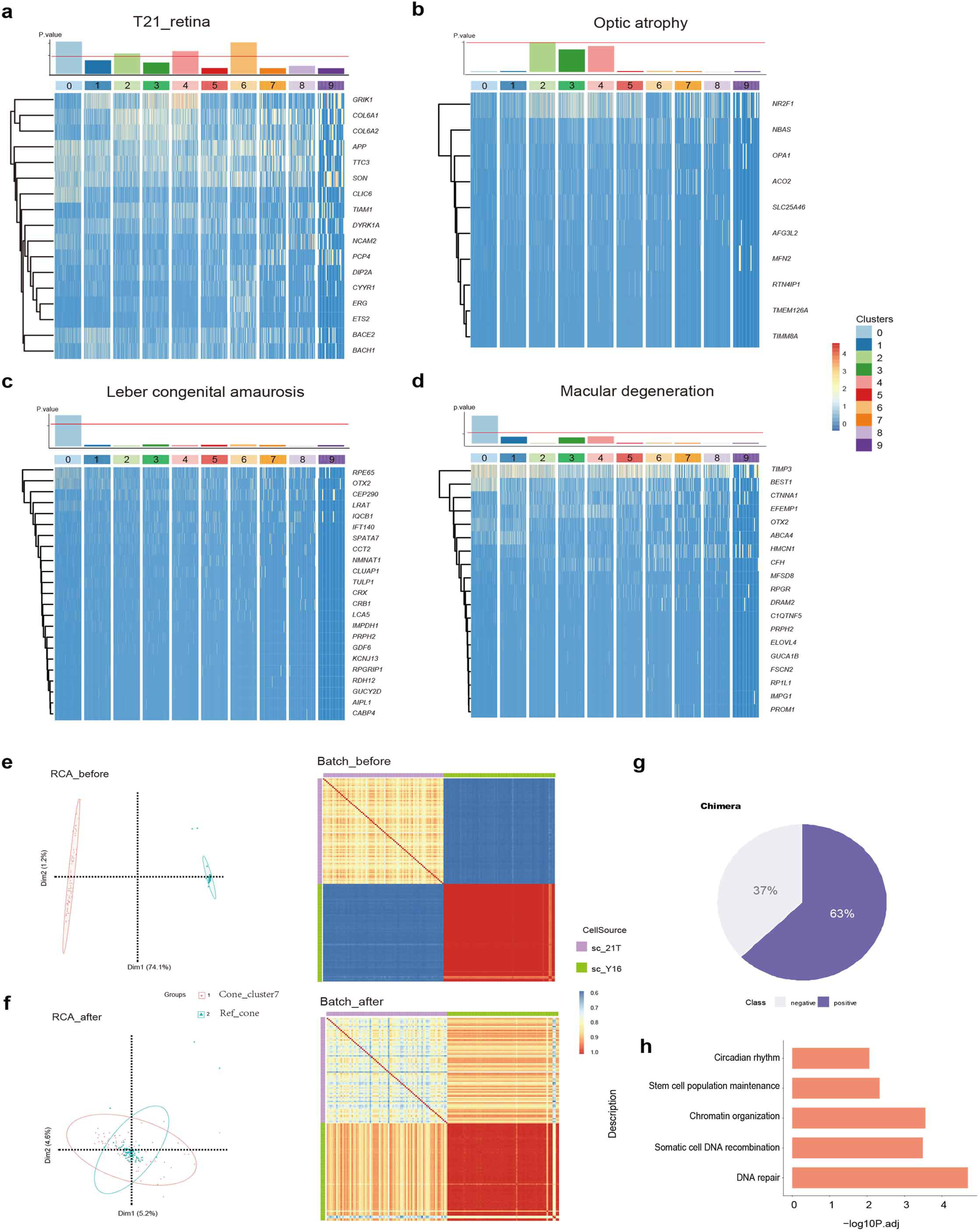
a. b. c. d. P values of enrichment of each cluster about trisomy 21 dosage genes and three retina diseases related genes and heatmaps of the corresponding significantly expressing genes. e. f. PCA plots and correlation heatmaps of before and after remove of batch effects respectively.. g. Prediction result of binary classification machining learning model. h. Enriched GO terms of variably expressed genes between 21 trisomy cells and normal cells

Leber congenital amaurosis and Macular degeneration associated genes were enriched in RPCs(C0) with the P value less than 0.05. Also, we noticed that C0 specifically expressed *RPE65, OTX2, LRAT* and *BEST1*^34,35^. While Optic atrophy associated genes was enriched in astrocyte(C2), where *NR2F1*^36^ and *AFG3L2*^37^ were highly expressed.

The obtained sample were chimera of chromosome 21, which was verified using karyotype detection. To explore the ratio of trisomy 21 cells and diploid 21 cells in our sample, we employed a training Random Forest classifier which fed with the normal counterpart single cell data^38^ and in-silico negative data then performed prediction on mixed status sample. The predict result manifested the proportion of cells with trisomy 21 in this sample was 0.63 (Figure 5)^39,40^, which consisted of 50 trisomy 21 cells and 39 diploid cells.

## Discussion

In this study, we conducted scRNA-seq of the retina from a trisomy 21 fetal to dissect the heterogeneity of retina, providing the first retinal cell atlas under T21 condition. We also studied the influence of redundant chromosome 21 in the process of retinogenesis through reconstructing pseudo-time trajectories.

The transcriptome of single cell of retinal tissue revealed significant heterogeneity within retina. The study identified subtypes of retinal progenitor cell exhibiting the potential of different retinal cell type commitments and characterized certain Müller glia cell with differentiating potency. Furthermore, we constructed a detailed communication network of Müller glia and other retinal neural cells, while the communication regulatory interactions of the subtypes of progenitor Müller glia were the most widely identified. As expected, the astrocytes were involved in the interactions of immune system regulation. As the mean of detected gene numbers of single cell transcripts sequencing is generally low, we validated the reliability of our data by identifying the detection of housekeeping genes. Overall, this study provides a data resource for the heterogeneity and development progression of trisomy 21 retina. The significant correlation between progenitor cells and trisomy 21 as well as retina disease provided orientation of the research about these retina diseases.

Finally, two retinal cell types, amacrine cells and horizontal cells, were found in another single cell human fetal study^11^ and several other mammalians retinal study^14^, but absent in a retinal organoids study^16^ speculatively due to the deficiency of sampled cells number. We assumed that the absence of these two cell types in our data might indicate the general delay of the development of trisomy 21 retina or the severe abnormity of highly chimera of trisomy 21. Further researches and experimental verifications are needed to test the above hypotheses.

## Materials and Methods

### Ethics statement

T21 human fetal retina collection and research was approved by Medical Ethics Committee of Shiyan Taihe Hospital (201813). The informed consent forms were designed under the ISSCR guidelines for fetal tissue donation and in strict observance of the legal and institutional ethical regulations for elective pregnancy termination from the patient after her decision to legally terminate her pregnancy but before the abortive procedure. All tissue samples used in this study were not previously involved in any other procedures. All protocols were in compliance with the ‘Interim Measures for the Administration of Human Genetic Resources’ administered by the Chinese Ministry of Health.

### Tissue sample collection and dissociation

We collected 23 weeks retina which from left eye of a T21 human fetal. Gestational age was measured in weeks from the first day of woman’s last menstrual cycle to the sample collecting date. T21 human fetal retina sample was collected into the Stroke-physiological saline solution. A rapid hemi-section was performed to remove the vitreous and the anterior. The retina was carefully dissected free from posterior eyecup and then flash frozen in liquid nitrogen. The tissue was provided with de-identified medical records including time and cause of death. The time between death and tissue collection was 3hrs.

### Single cell cDNA library preparation and high-throughput sequencing

Flash-frozen tissue was homogenized in 2mL ice-cold Lysis buffer (10 mM Tris-HCl, pH 7.4, 10 mM NaCl, 3 mM MgCl2, 0.1% NP40, protease inhibitors) then dounced in an RNase-free 2ml glass dounce (D8938-1SET SIGMA) 15x with a loose pestle and 15x with tight pestle on ice. Transfer homogenization through 40 µm filter (352340 BD) and removed the block mass. The cell filtrate was proceeded to Density Gradient Centrifugation. Mixed 400 µL cell filtrate with 400 µL 50% Iodixanol Solution in 2mL lo-Bind tubes (Z666556-250EA SIGMA). Carefully layer 29% Iodixanol Solution and 35% Iodixanol Solution to the bottom of tube. Centrifuging at 3,000 g, 4°C for 30min. Nuclei were resuspended in ice-cold 1 x PBS (10010-031 GIBCO) containing 0.04% BSA and spin down at 500g for 5min. Discarded supernatant and then using regular-bore pipette tip gently pipette the cell pellet in 50 μL ice-cold 1 x PBS containing 0.04% BSA and 0.2U/µl RNase Inhibitor. Determined the nuclei concentration using a Hemocytometer (101010 QIUJING). Then, loaded on a Chromium Single Cell Controller (10x Genomics) to generate single-cell Gel Bead-In-EMulsions (GEMs) by using Single Cell 3’ Library and Gel Bead Kit V2 (120237 10x Genomics). Captured cells released RNA and barcoded in individual GEMs. Following manufacturer’s instructions (120237 10x Genomics) library was generated from the donor sample. Indexed library was converted by MGIEasy Lib Trans Kit (1000004155, MGI) then sequenced on the MGISEQ 2000 (MGI) platform with pair-end 26bp+100bp+8bp (PE26+100+8).

### Pre-processing and quality control of scRNA-seq data

We first used Cell Ranger 2.0.0 (10X Genomics) to process raw sequencing data and then applied Seurat (10.1038/nbt.3192) for downstream analysis. Before we start downstream analysis, we focus on four filtering metrics to guarantee the reliability of our data. (1) We filter out genes that are detected in less than 0.1% of total cell number to guarantee the reliability of each gene; (2) We filter out cells whose percentage of expressed mitochondrial genes is greater than 10%; (3) We also filter out cells whose UMI counts is either less than or greater than one IQR distance outer of the quartiles of UMI counts to filter out cells; Finally, we use the house keeping genes from Protein Alta(http://www.proteinatlas.org/) to verify the reliability of our data.

### Analysis of heterogeneity in each tissue and cell line

The heterogeneity of the retina sample was determined using Seurat R package (10.1038/nbt.3192). Then we determined significant PCs using the JackStrawPlot function. The top twelve PCs were used for cluster identification with resolution 1.0 using k-Nearest Neighbor (KNN) algorithm and visualization using t-Distributed Stochastic Neighbor Embedding (tSNE) algorithm. Cell type were assigned by the expression of known cell-type markers and functional enrichment analysis. The FindAllMarkers function in Seurat was used to identify marker genes for each cluster using default parameters. Removal of cell cycle effects in clustering and cell cycle analysis. We collected 43 genes and 54 genes related to S phase and G2/M phase respectively (10.1126/science.aad0501; 10.1101/gr.192237.115). For clustering, each cell was assigned a score to describe its cell cycle state by CellCycleScoring function in Seurat according to the expression of these genes. Subsequently, the cell cycle effect was regressed out based on the scores, leading to a more accurate clustering result. For cell cycle analysis, cells were determined to be quiescent (G1 stage) if their S score < 0 and G2/M score < 0; otherwise, they were deemed proliferative. In addition, proliferative cells were designated G2/M if their G2/M score > S score, whereas cells were designated S if their S score > G2/M score.

### GO term and KEGG pathway enrichment analysis

Lists of genes were analyzed using clusterProfiler R package (10.1089/omi.2011.0118) and the BH method was used for multiple test correction. GO terms with a P value less than 0.05 and KEGG term with a P value less than 0.05 were considered as significantly enriched. GO terms enrichment analysis of target genes of TFs used *Metascape (*http://metascape.org/gp/index.html), which is flexible for gene multiple functional analysis.

### Construction of trajectory using variable genes

Monocle (10.1038/nbt.2859) ordering was conducted for constructing single cell pseudo-time of retinal cells.using highly variable genes identified by Monocle to sort cells in pseudo-time order with default parameters.. “DDRTree” was applied to reduce dimensional space, and the minimum spanning tree on cells was plotted by the visualization functions “plot_cell_trajectory” or “plot_complex_cell_trajectory”. BEAM tests were performed on the first branch points of the cell lineage using all default parameters. Plot_genes_branched_pseudotime function was performed to plot a couple of genes for each lineage.

### Regulatory network construction

We downloaded human TF list from AnimalTFDB (10.1093/nar/gkr965) as a TF reference and extracted TFs in marker genes list of each cluster to construct the regulatory network. The extracted TFs were submitted to STRING database (10.1093/nar/gkw937) to infer regulatory networks based on known interaction relationships (supported by data from curated databases, experiments and text-mining). TFs without any interactions with other proteins were removed from the network.

### Construction of cellular communication network

The ligand-receptor interaction relationships were downloaded from the database, IUPHAR/BPS Guide to PHARMACOLOGY (10.1093/nar/gkx1121), and the Database of Ligand-Receptor Partners (DLRP) (10.1101/gr.207597.116; 10.1093/nar/gkh086). The average expression level of UMI number of 1 was used as a threshold. Ligands and receptors above this threshold were considered as expressed in the corresponding cluster. The R package Circlize (10.1093/bioinformatics/btu393) was used to visualize the interactions.

### Construction of cross-tissue and cross cell type correlation network

To reduce noise, we averaged the expression of every 30 cells within cluster and then calculated the pairwise Pearson correlation between two dots based on their average expression profiles. Inter-dots relationship will be shown if their Pearson correlation is greater than 0.95. This correlation network is generated using Cytoscape (10.1101/gr.1239303).

### Enriched ontology clusters

We first identified all statistically enriched terms, accumulative hypergeometric p-values and enrichment factors were calculated and used for filtering. Remaining significant terms were then hierarchically clustered into a tree based on Kappa-statistical similarities among their gene memberships. Then 0.3 kappa score was applied as the threshold to cast the tree into term clusters (stable4). We then selected a subset of representative terms from this cluster and convert them into a network layout. More specifically, each term is represented by a circle node, where its size is proportional to the number of input genes fall into that term, and its color represent its cluster identity. The network is visualized with Cytoscape (v3.1.2) with “force-directed” layout and with edge bundled for clarity. One term from each cluster is selected to have its term description shown as label.

### Protein-protein interaction network

MCODE algorithm was then applied to this network to identify neighborhoods where proteins are densely connected. Each MCODE network is assigned a unique color. GO enrichment analysis was applied to each MCODE network to assign “meanings” to the network component.

### Binary classification of trisomy 21 and normal cell by machining learning

#### 1. Standardization

Since the difference between read counts based on the third-party data and UMI counts based on our experiments, standardization was employed to unify the distributions in the two different sources. Theoretically the gene expression level is as a continuous random variable subject to the normal distribution. We employ centralization and variance normalization to normalize both datasets on the basis of read counts and UMI counts both considered as finitely sampling from the intrinsic gene expression and converged to the normal distribution.

#### 2. Batch effects normalization

In analysis of single-cell mRNA-seq data, batch effects give rise to non-negligible deviation. Map points shown as two clusters from two sources present significant batch effects including library preparations, read-mapping methods and etc. This effect performs the first components beyond intrinsic biological differentiate. Batch effects were adjusted by sva package in R.

#### 3. Generate in-silico negative samples

For discrimination of normal cells from mixed samples, we generated in-silico negative samples of the same amount as normal cells based on some prior knowledge about abnormal expression of trisomy 21 cells.

The related experiment was divided into five steps:

1. Count the amounts of different genes appearing in all datasets as features to construct the feature space.
2. Generate the preliminary sample vector in the feature space from the normal distribution N(1, 0.01).
3. Sample from a given set of highly expressed genes (highly expressed in trisomy 21) to obtain a subset of highly expressed genes with random numbers and positions.
4. Up-regulate the gene expression level at the corresponding position of the preliminary sample vector.
5. Sample vector center and variance normalization.

#### 4. Data sharding & model fitting

In-silico samples were mixed into the real-world datasets after all normalization completed and during this mixture one special label in normal or abnormal was tagged to each sample, which helped subsequently training classifier. Then eighty percent of mixed data was split to feed Random Forest and twenty percent left was to evaluate the model. Random Forest was fitted with default parameters and dumped for permanent access with 0.97 OOB score and 0.98 AUC score on test dataset.

#### 5. Prediction

The dumped model performed predictions on our datasets where normal cells and abnormal cells had not been distinguished. The predictions were mapped into t-SNE plot of dimensional reduction of origin data with no standardization and normalization.

#### 6. Optimizing

Lack of sufficient prior knowledge, the distribution of the in-silico simulated samples have great differences with the trisomy 21, which makes it unreasonable to decide the abnormal cell predicted by Random Forest as the trisomy 21. There are too many unwanted variations between the two data so that it is hard to control moderate adjustment. Over-correction of batch effects possibly eliminates the intrinsic biological difference. PCA pre-processing can be alleviated before classification. Additionally, too few cells sequenced and too many genes measured increase risk of over-fitting, which can be alleviated by dimensional reduction of PCA or other tools.

## Acknowledgements

This work was supported by NSFC 81770925, 81790641; the Non-profit Central Research Institute Fund of Chinese Academy of Medical Sciences 2018PT32019.

## Author contributions

JW, DC, XS, FC, FH conceived the project. JZ, ZH, SX, GG, PX, FG, DW, DZ coordinated and collected the human donor retinas, XD, XL, QF, JZ, JZ, WL performed scRNA-seq data analysis. scRNA-seq library generation and protocol optimization performed by ZX, LL, CC. ZX, SW generated the scRNA-seq data (with help from JX). YC, XD developed the binary classification model. JW, XD, FH wrote the manuscript with input from all authors. DC, XS, FC, SJ, JL, YG revised the manuscript.

---

**Figure s1:**
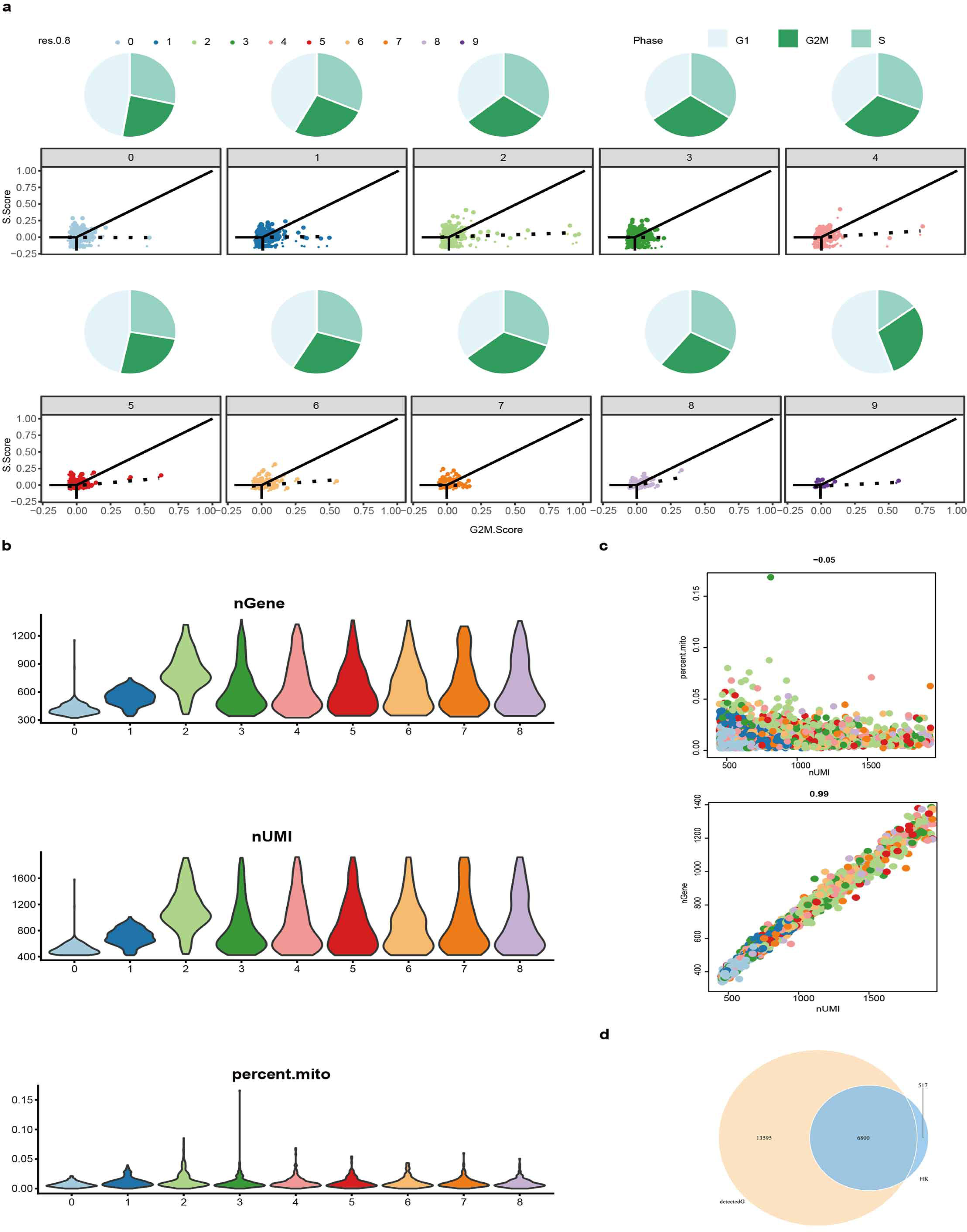
a: Visualization of cell cycles of clusters by pie chart and scatter plot; b: Presentation of the gene expression number(above), UMI number(intermediate) and the mitochondrial genes proportion (below) in clusters by violin plots; c: Correlations of the UMI and mitochondrial genes(above) as well as the UMI and gene(below). d: Venn plot shows housekeeping gene expression in our data set.

**Figure s2:**
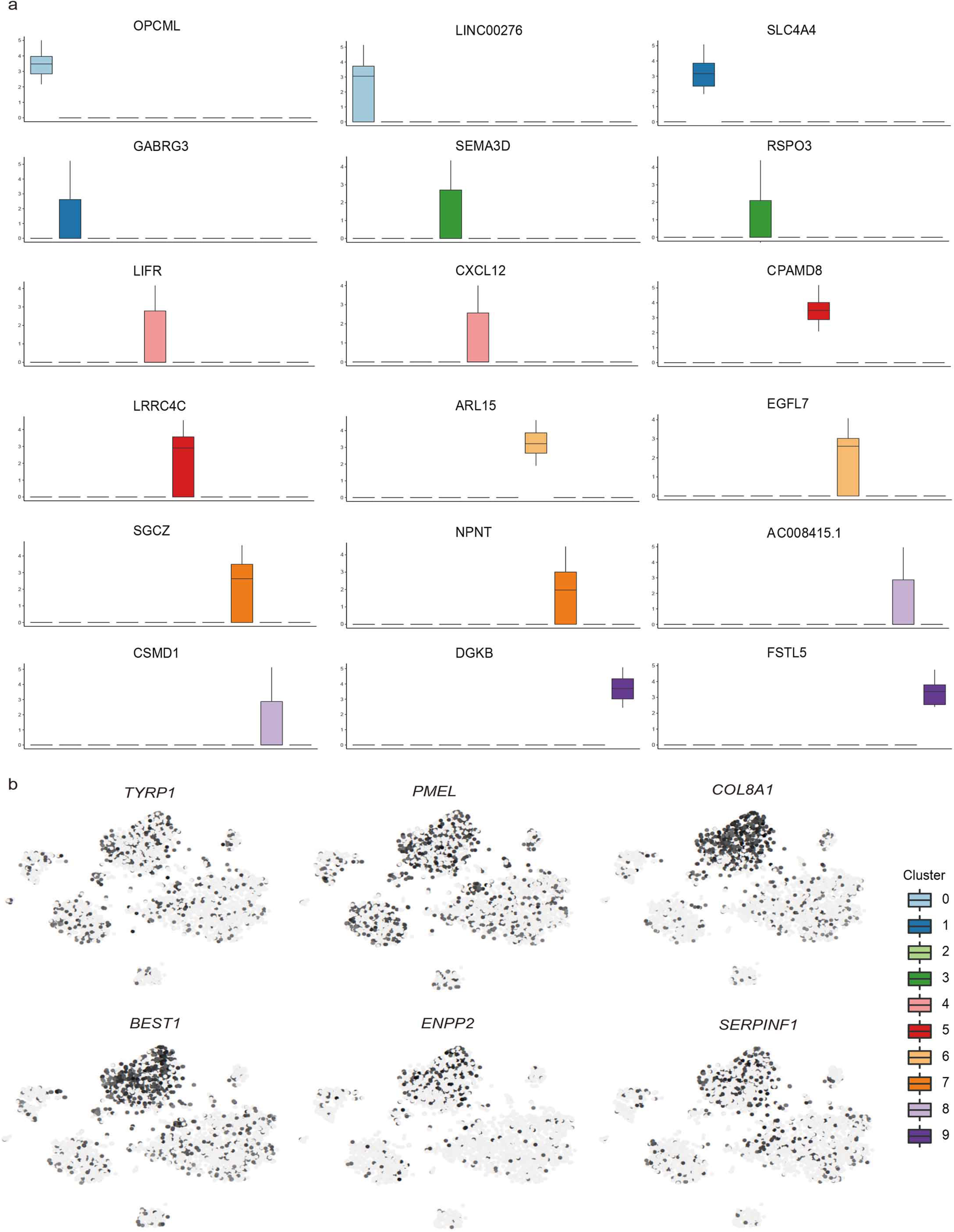
a: Box plot of genes specifically expressed in clusters; b: Expression of melanin markers in clusters.

**Figure s3:**
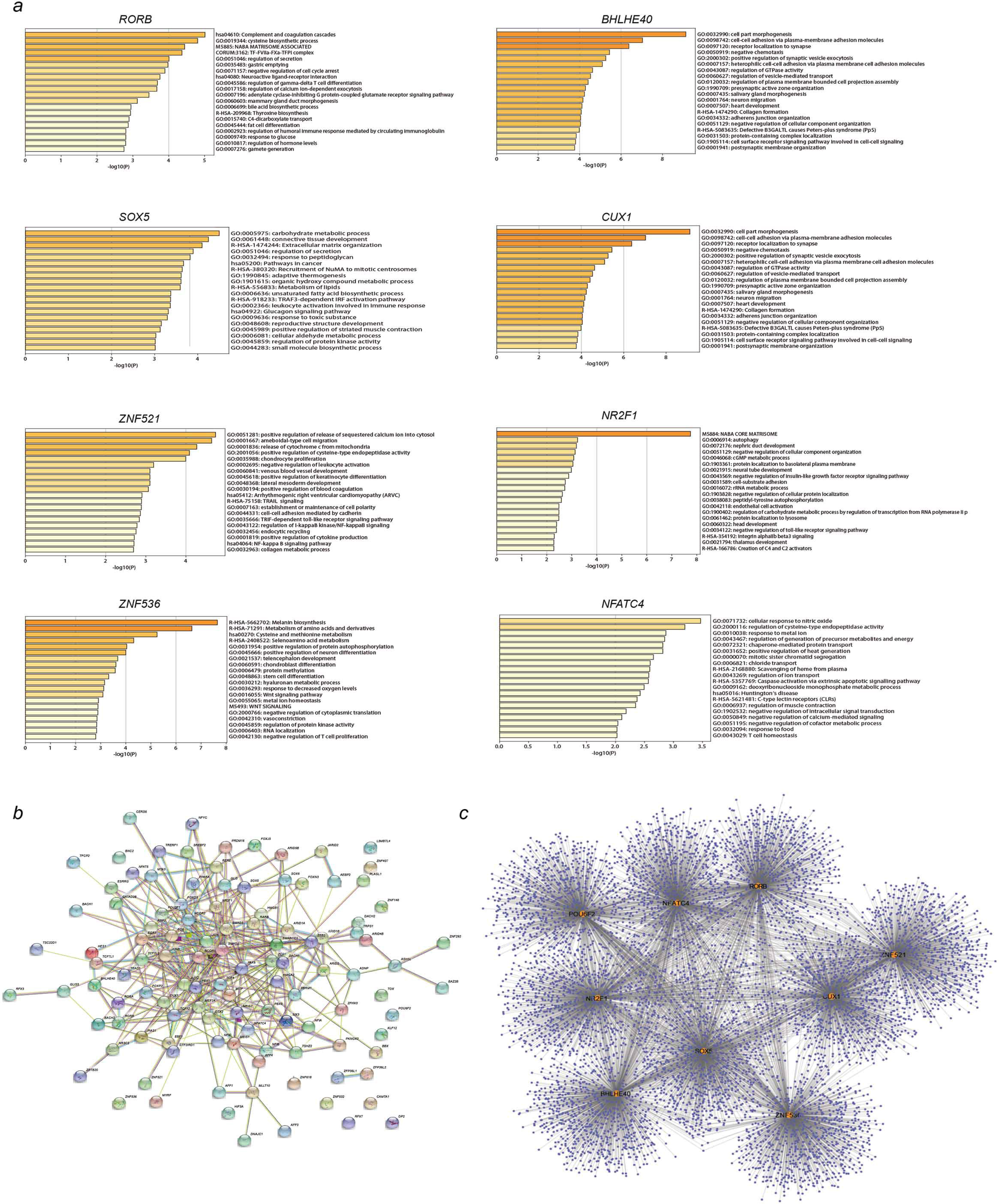
a: Enriched GO terms of top1000 predicted target genes of variably expressed TFs along the trajectory. b: Gene regulatory network of top100 variably expressed TFs. c: Regulatory network of top1000 predicted target genes of selected variably expressed TFs.

**Figure s4:**
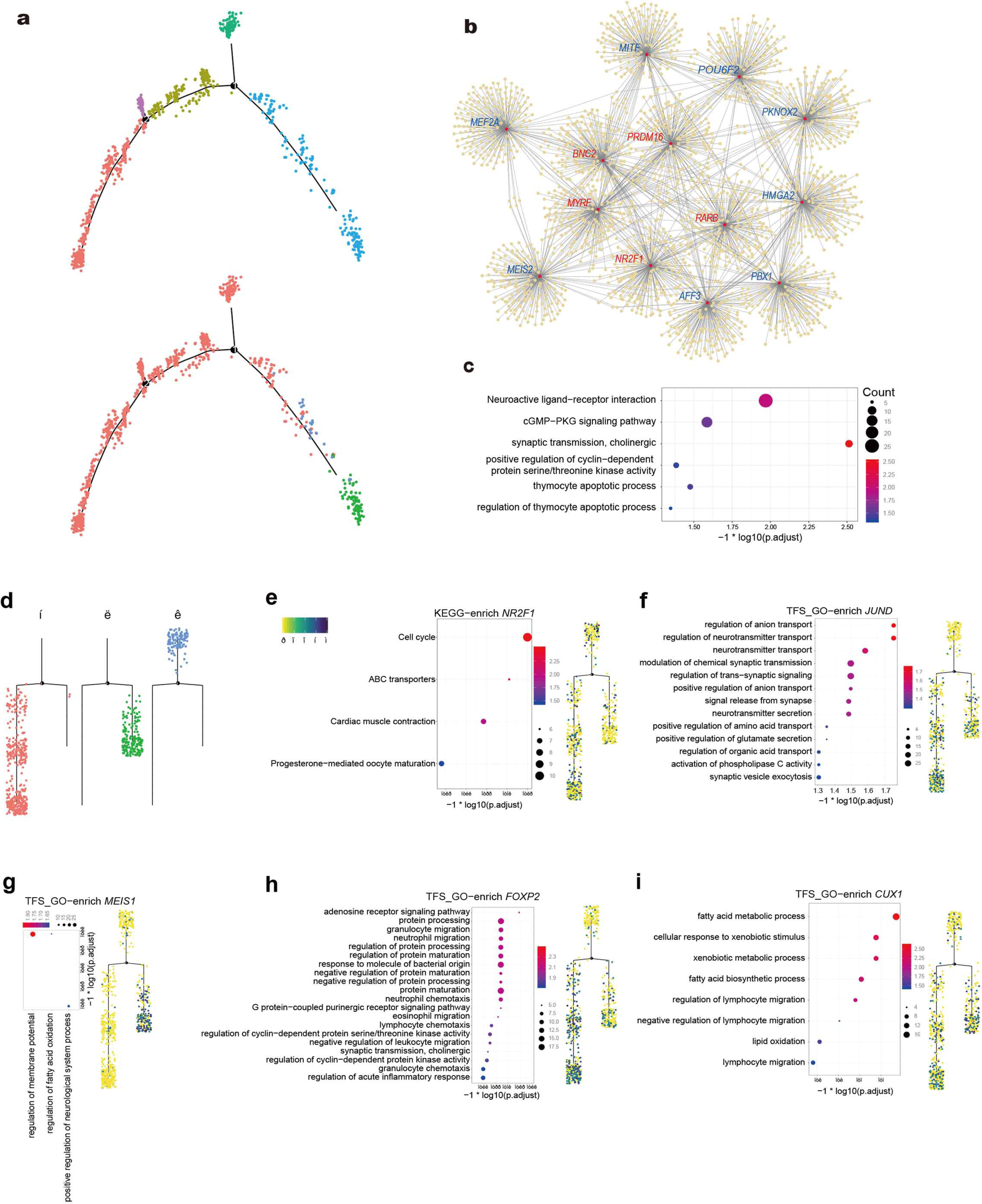
a, d: Analysis of development trajectories of cone cells and the muller cells. a: according to cell differentiation status(above) and cell subpopulations(below); d: differentiation trajectories of the three subpopulations of muller cells. b: Regulatory network of the cone cells c: Go term enrichment analysis of the differentially expressing TFs in cone subtypes. e - i: Enrichment analysis of specific transcriptional regulators in different differentiated branches

**Figure s5:**
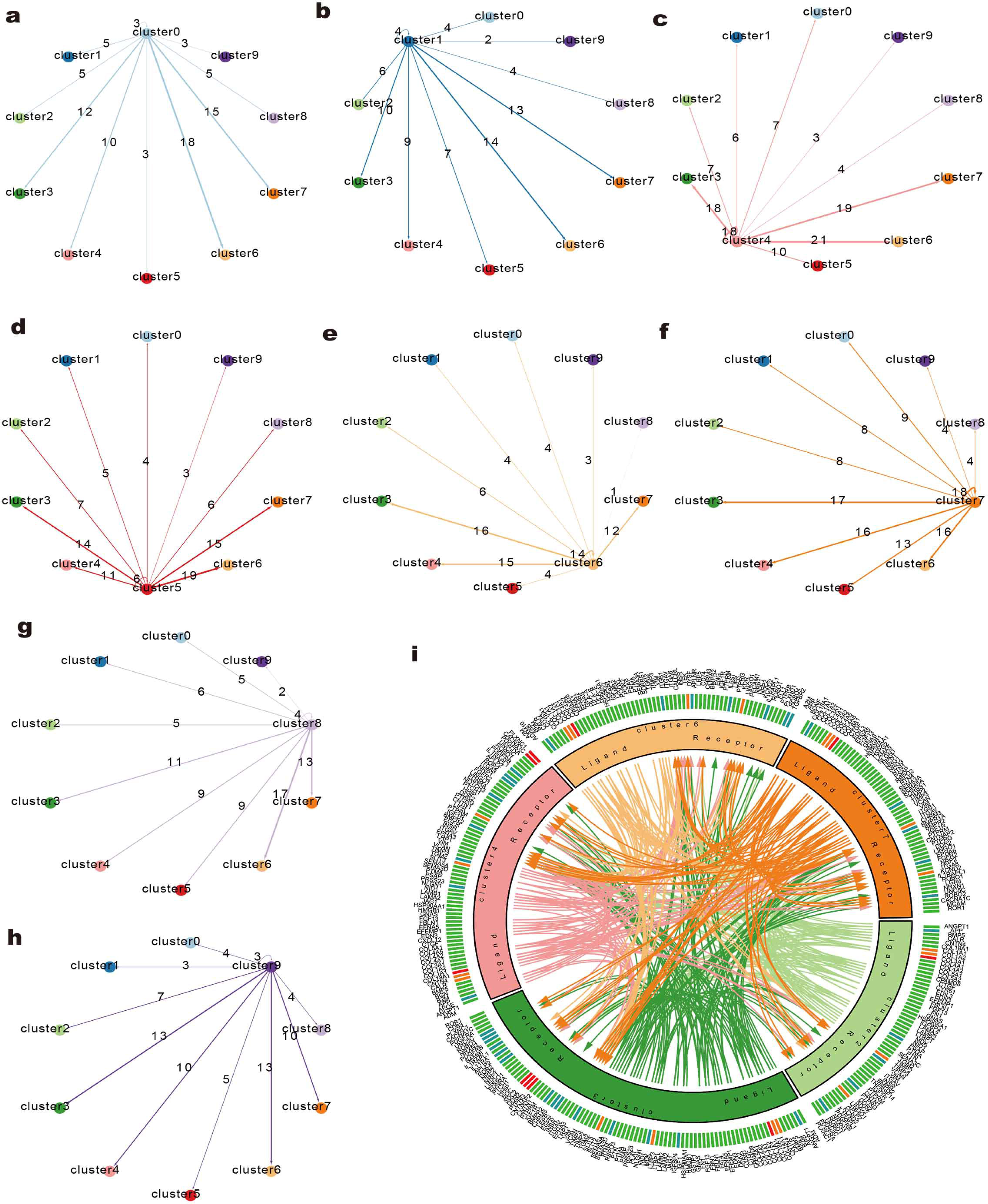
a-h: The interaction between clusters characterized by the number of ligands and receptors; i: The detail visualization of the interaction between clusters 2/3/4/6/7.

**Figure s6:**
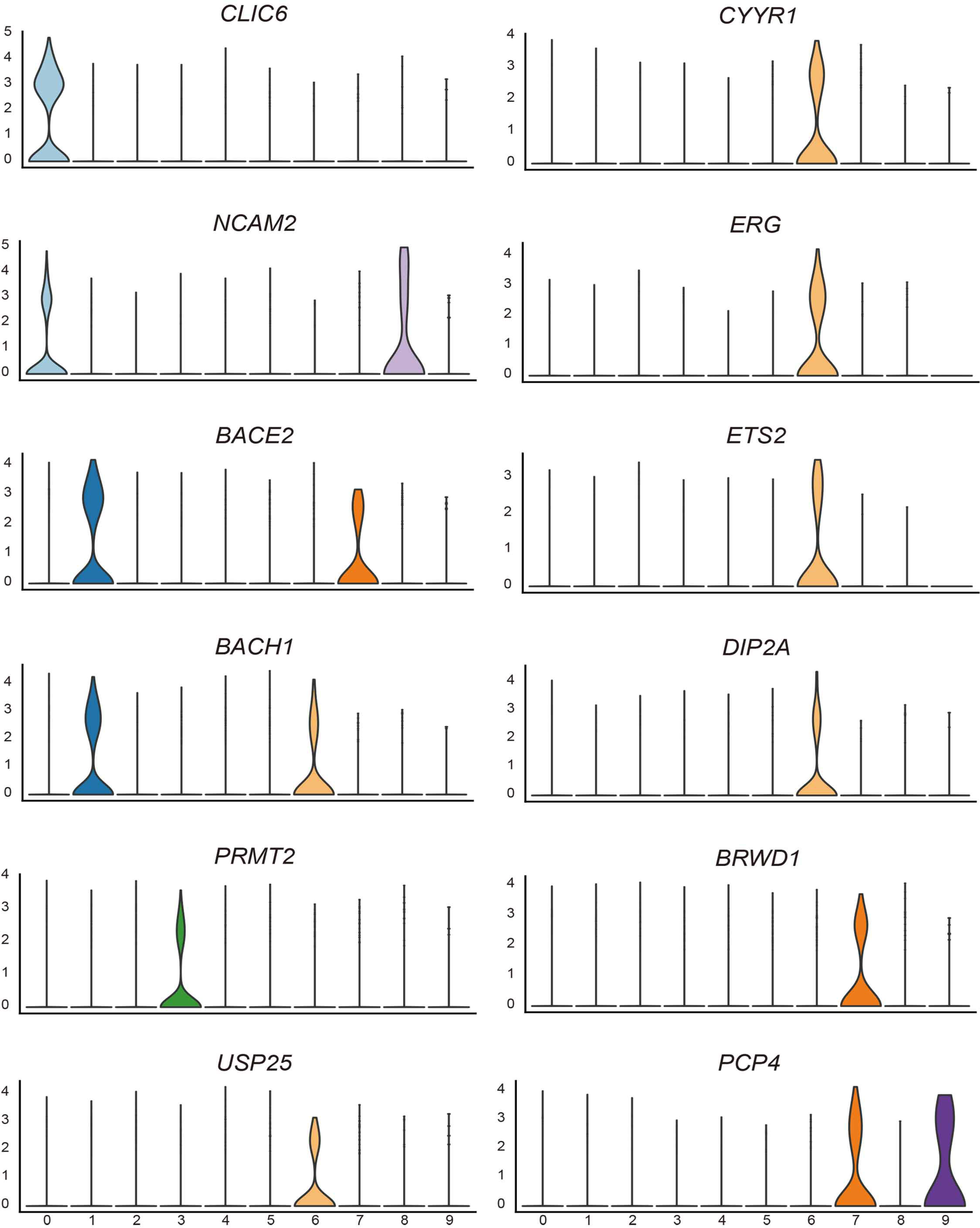
Violin plots of genes on chromosome 21 specifically expressed in clusters

